# Plant growth regulators interact with elevated temperature to alter heat stress signaling via the Unfolded Protein Response in maize

**DOI:** 10.1101/532796

**Authors:** Elena M. Neill, Michael C. R. Byrd, Thomas Billman, Federica Brandizzi, Ann E. Stapleton

**Affiliations:** University of North Carolina Wilmington, Department of Biology and Marine Biology, Department of Mathematics and Statistics, Wilmington, NC, USA; Michigan State University, Department of Plant Biology, East Lansing, MI, USA; North Carolina Department of Health and Human Services, State Laboratory of Public Health, Raleigh, NC, USA; Blue Cross and Blue Shield of North Carolina, Underwriting Division, Durham, NC, USA

## Abstract

Plants are increasingly exposed to high temperatures, which can cause accumulation of unfolded protein in the endoplasmic reticulum (ER). This condition, known as ER stress, evokes the unfolded protein response (UPR), a cytoprotective signaling pathway. One important branch of the UPR is regulated by splicing of bZIP60 mRNA by the IRE1 stress sensor. There is increasing evidence that commercial plant growth regulators may protect against abiotic stressors including heat stress and drought, but there is very little mechanistic information about these effects or about the regulatory pathways involved. We evaluated evidence in the B73 Zea mays inbred for differences in the activity of the UPR between permissive and elevated temperature in conjunction with plant growth regulator application. Treatment with elevated temperature and plant growth regulators increased UPR activation, as assessed by an increase in splicing of the mRNA of the IRE1 target bZIP60 following paclobutrazol treatment. We propose that plant growth regulator treatment induces bZIP60 mRNA splicing which ‘primes’ plants for rapid adaptive response to subsequent endoplasmic reticulum-stress inducing conditions.

## Introduction

Improving crop production, especially maize (Zea mays L.) production, has become a topic of increasing interest globally due to climate change, population growth and alternative crop uses such as biofuels and plastics^1^. Global patterns in cereal grain production for maize, rice, and wheat have been uneven in that certain areas of the world have experienced increases in production while other areas have not. To meet the food demands of an increasing global population, it will necessary to double grain production by 2050^2^. While the scientific and agricultural community has been able to increase grain production in past years by breeding hybrid crops that are more resistant to both biotic and abiotic stressors, these challenges may be more difficult to meet in the face of climate change. The problem is multifaceted; climate change and the ability of farmers to predict environmental trends are both key for crop yield. Estimates for maize suggest that exposure to temperatures above 29 degrees C for even one day decreases overall yield by half a percent^3^. Much is already known about the mechanisms which plants use to respond to heat stress, but continuing research will allow geneticists to further explore treatments which may influence stress response in plants, and forestall the events that reduce crop yield^4^.

Heat stress disrupts sensitive molecular processes such as the folding of proteins in the endoplasmic reticulum (ER) of plant cells. The accumulation of misfolded proteins in the ER is a potentially toxic condition defined as ER stress. ER stress elicits the unfolded protein response (UPR), a common adaptive response to mitigate stress and to protect from further stress^5^. However, in the face of persistent stress the UPR transitions from adaptive responses to programmed cell death^6^. During the UPR, the ER signals the nucleus to upregulate the expression of stress response genes. Two UPR signaling arms exist in plants^7^. One arm involves the ER membrane-associated transcription factor bZIP28, which is mobilized from the ER to nucleus in response to stress. The other arm involves the ER membrane-associated kinase and ribonuclease IRE1, which splices the mRNA of bZIP60, a potent transcription factor, in response to stress^8^. The splicing is critical because it results in the activation of bZIP60, which is translocated to the nucleus where it modulates the expression of UPR target genes in a manner partially overlapping with bZIP28^9, 10^ Because the IRE-specific splicing of bZIP60 mRNA is a key step in the response to ER stress, it can be used as a reliable indicator for the induction of the UPR. The UPR is activated in dicots and monocots in response to heat, as supported by an enhancement of bZIP60 splicing in Arabidopsis and Brachypodium seedlings upon exposure to elevated temperature^7^. It has recently been suggested that the PIF4 family of transcription factors control heat and phytohormone responses, possibly in a manner that is independent from other signaling pathways such as those regulated by the UPR^11^.

There is substantial information thermosensing pathways, which are currently grouped into PIF4 and non-PIF4 subsets^11^. The Li (2018)review summarized regulatory interactions into two groups, first those that include PIF4 in heat and phytohormone responses and a second set of interactions which includes the CPR, UPR, and MAPK responses. The Li (2018) review also summarizes how paclobutrazol inhibits thermo-responsive hypocotyl growth via DELLA in the PIF4 regulatory network. We thus have some information on phytohormone involvement in PIF4 pathways, but we don’t know if hormones are involved in non-PIF4 pathways. Thus, testing for hormone involvement in the UPR would begin to define the connection between non-PIF4 and PIF4-related heat responses, which was identified by Li (2018) as a key open question.

Plant growth regulators are commercially available chemicals that are used abundantly in crop production to manage disease, promote growth, and increase yield^12^. While there are approximately 40 plant growth regulator compounds with a variety of active ingredients, all plant growth regulators share the common function of regulating intrinsic hormone levels within plants by modulating signaling within various hormone transduction pathways^12^. Plant growth regulators can be better described as “plant bioregulators” and can be further subdivided into “plant biostimulants” and “plant retardants” as well as into five other groups: (1) auxins, (2) gibberellins, (3) cytokinins, (4) abscisic acid, and (5) ethylene^12^. Gibberellins, auxins, and brassinosteroids contribute to plant height and organ size, but the effects of these hormones are not additive^13^. Agronomic use of these plant growth regulators is focused on amelioration of biotic and abiotic stress in agronomic applications^14–16^ For future discovery and eventual rational design of regulators we will require mechanistic information about mode of action and interactions of these compounds.

In this study we investigated the role of four plant growth regulators – propiconazole, uniconazole, paclobutrazol, and Quilt Xcel – on heat induction of the UPR in maize. Propiconazole, uniconazole, and paclobutrazol belong to the triazole family of compounds and affect gibberellic acid, abscisic acid, and brassinosteroid biosynthesis^12^. Triazoles directly impact gibberellic acid synthesis which retards growth and lead to the formation of more compact plants with less stem elongation^17^. Compact plants have advantages such as less energy expenditure towards shoot growth, which allows the plant to put in more energy towards flowering, reproduction, and survival^17^. Paclobutrazol has recently been shown to reduce lodging and increase stem strength in maize, along with reducing plant height^18, 19^; low levels of uniconazole application increased abscisic acid and decreased gibberellic acid in both foliar and leaf treatments in maize kernels^20^ and reduced plant lodging^21^.

Quilt Xcel includes azoxystrobin, which has a different mode of action than the triazole compounds, instead working as a quinone outside inhibitor which acts directly on the quinol outer binding site of the cytochrome bc1 complex^22^. The cytochrome bc1 complex is involved in electron transport for ATP production and in the regulation of mitigating free radical damage that may cause cellular damage due to reactive oxygen species arising from the formation of a superoxide ion. It has been observed that despite the presence of abiotic and biotic stressors, strobilurin compounds such as Quilt Xcel may serve to increase crop yield, but the effect of this treatment has not been extensively evaluated at a mechanistic level in the public scientific literature. The active ingredients in Quilt Xcel are 13.5 percent azoxystrobin and 11.7 percent propiconazole (Syngenta Inc, Basel, Switzerland).

The goal of this study was to investigate the connections between plant growth regulators and heat induction of the UPR. Previous studies have noted that either heat stress or the action of ER stress agents, such as tunicamycin or dithiotheitol, increases the splicing of bZIP60 mRNA, a biomarker for the UPR^23^. Thus our primary objective was to determine if the preemptive application of plant growth regulator treatments on maize seedlings influences the splicing bZIP60 mRNA in response to heat stress. In addition, we analyzed the effects of plant growth regulator treatments on plant growth to determine if there was a significant interaction between temperature and plant growth regulator treatment. We found that there were significant interactions between some plant growth regulator treatments and temperature for phenotypic traits and a significant increase in the UPR from the untreated compared to plant growth regulator treatments. Temperature interactions with plant growth regulator treatment were present in some combinations of plant growth regulator and heat, though the effect of heat on the UPR contributed less to overall ER-splicing effects than did plant growth regulator treatments.

## Methods

### Seed Stocks

The Zea mays B73 inbred genotype seed was provided by the Maize Co-op (http://maizecoop.cropsci.uiuc.edu/) and seeds stocks were increased using standard maize nursery self-pollination methods at the Central Crops agricultural experiment station, Clayton, NC (http://www.ncagr.gov/research/ccrs.htm). Seed genotypes were verified each season using SSR marker comparison to the standards in the MaizeGDb listing, https://www.maizegdb.org/data_{_}center/ssr.

### Experimental Design and Growth Conditions

Maize B73 seeds were grown at the North Carolina State Phytotron facility in two climate controlled growth chambers. Seeds were planted in a 50:50 soil/gravel mixture on November 5, 2016. To ensure uniform growing conditions, the depth at which each seed was covered in soil did not exceed three times the seed’s length. The experiment consisted of a randomized pair block design as shown in 1. Twenty plants were randomly distributed within two temperature-controlled chambers, one normal temperature chamber and one heat stress chamber. Within each chamber, five plants were designated to receive one of five potential treatments: no hormone, paclobutrazol, uniconazole, propiconazole, or Quilt Xcel.

Conditions in each chamber were held constant through the V4 plant growth stage to simulate growing conditions found in at the start of a typical growing season in the primary Corn Belt of the United States. The growing temperature during the day and night was 28 degrees C and 21 degrees C respectively. The distribution of plants within each chamber was randomized in order to ensure that different locations within the chamber would not lead to confounding effects. Temperature, light, watering conditions, and humidity levels were also maintained on a regular schedule throughout the entirety of the experiment according to the NC State Phytotron Procedural Manual. Cool white fluorescent (1500 ma) and incandescent lights remained on from 7:00 AM until 9:00 PM and off during the remainder of the day. Relative humidity in the C-chambers was maintained between 60-70 percent. Plants were watered with deionized water in the morning and a nutrient mixture in the afternoon (Phytotron Procedural Manual, 2009). These conditions were kept constant throughout the two-week growing period.

### Hormone Treatment and Heat Ramp

Each plant growth regulator treatment was diluted to a 50 ppm concentration from the commercially available products. Uniconazole (Sumagic,Valent, Inc.) was diluted from a 443 ppm stock to a 50 ppm stock. Piccolo Ornamental plant growth regulator (Fine America, Inc.) contained the main active ingredient paclobutrazol, which was diluted from the 143,000 ppm stock concentration to 50 ppm. Propiconazole14.3 (Quali-Pro, Inc.) was diluted from a starting stock concentration of 550 ppm to 50 ppm. Quilt Xcel (Syngenta Inc.) was prepared from a 252,000 ppm stock concentration to a 50 ppm 100 mL solution. On November 19th, 2016, 10 mL of each 50 ppm plant growth regulator was administered to each respective replicate series within each chamber.

One week after growth regulator application the heat ramp in growth chamber 10 was implemented. The ramp conditions were set as following: for temperature ramp 1 the day temperature was adjusted from 28 degrees Celsius to 33 degrees Celsius, while the night temperature was adjusted from 21 degrees Celsius to 23 degrees Celsius. These conditions remained stable for 24 hours before a second heat ramp was implemented. The day temperature was adjusted from 33 degrees Celsius to 38 degrees Celsius. The night temperature was increased from 23 degrees Celsius to 25 degrees Celsius. These conditions were held constant for a total of 60 hours until sampling was completed.

### Leaf Tissue Sampling and Collection of Phenotypic Data

Leaf tissues samples were collected from the longest leaf of each plant (leaf 4) using a pair of steel scissors which were bonded together in order to produce equal-sized strips of leaf tissues. These leaf tissue samples were placed in labeled centrifuge tubes containing 500 microliters of RNAlater solution (ThermoFisher Scientific, Inc., Atlanta, GA) and the filled tube promptly submerged in liquid nitrogen in order to prevent RNA degradation.

Phenotypic data including plant height, above-ground leaf and stem mass, and root mass was collected immediately after leaf tissue sampling. Height data was obtained by measuring the height in centimeters from the base of the plant to the most recently developed leaf ligule. Root mass was determined by separating the wet root mass from the stem and leaf tissues for each plant and allowing complete drying with no continued change in dry weight before recording the final dry mass of roots and above-ground leaf and stem tissues.

### RNA Extraction

The Qiagen RNeasy Plant Mini Kit was used to extract RNA from leaf tissue samples. Procedures in the fourth edition of the RNeasy quick start protocol were followed (Qiagen, Inc, Germantown, MD USA), with dithiothreitol used to prepare the RLT buffer. Since leaf tissue samples were stored in RNAlater solution to aid in preservation, the solution was quickly drained before the leaf segment was deposited in a 14 mL polystyrene round-bottom tube filled with enough liquid nitrogen to ensure samples would not thaw. The tissue was ground thoroughly with a glass pestle and RLT buffer applied. After extraction the RNA samples were stored at −80 degrees C until they were used for realtime qPCR.

### Realtime qPCR

The Bzip60exon1_2 and Across_IRE_Exon gene sequences from B73 genome v4 were obtained from the Maize GDB database (sequence details are available in Supplemental Data). Each gene sequence was copied into the Integrated DNA Technology (IDT, Inc.) PrimerQuest tool and the most optimal sequence was chosen according to assay optimization guidelines outlined on the IDT website. Probes specific to the small IRE-spliced intron were tested but failed quality control; we thus used a dual-assay design with the ratio of measurements for each sample. This ratio analysis only measures relative splicing levels; the amount of message cannot be compared across samples.

All realtime PCR was carried out by ARQGenetics, LLC (Bastrop, TX) using a BioRadCFX384 instrument (BioRad, Inc, Hercules, CA, USA). Calibration reactions with gBlocks serial dilutions and positive and negative controls were included on the same reaction plate as the samples Table 1. All reactions were carried out using BioRad standard operating procedures, with calibration of the CFX384 realtime instrument as specified by the manufacturer. RNA samples were reverse-transcribed and amplified using the BioRad Universal OneStep kit, with two microliters of template per reaction. Raw fluorescence count data were exported using the CFXManager software for the data analysis.

**Table 1.**
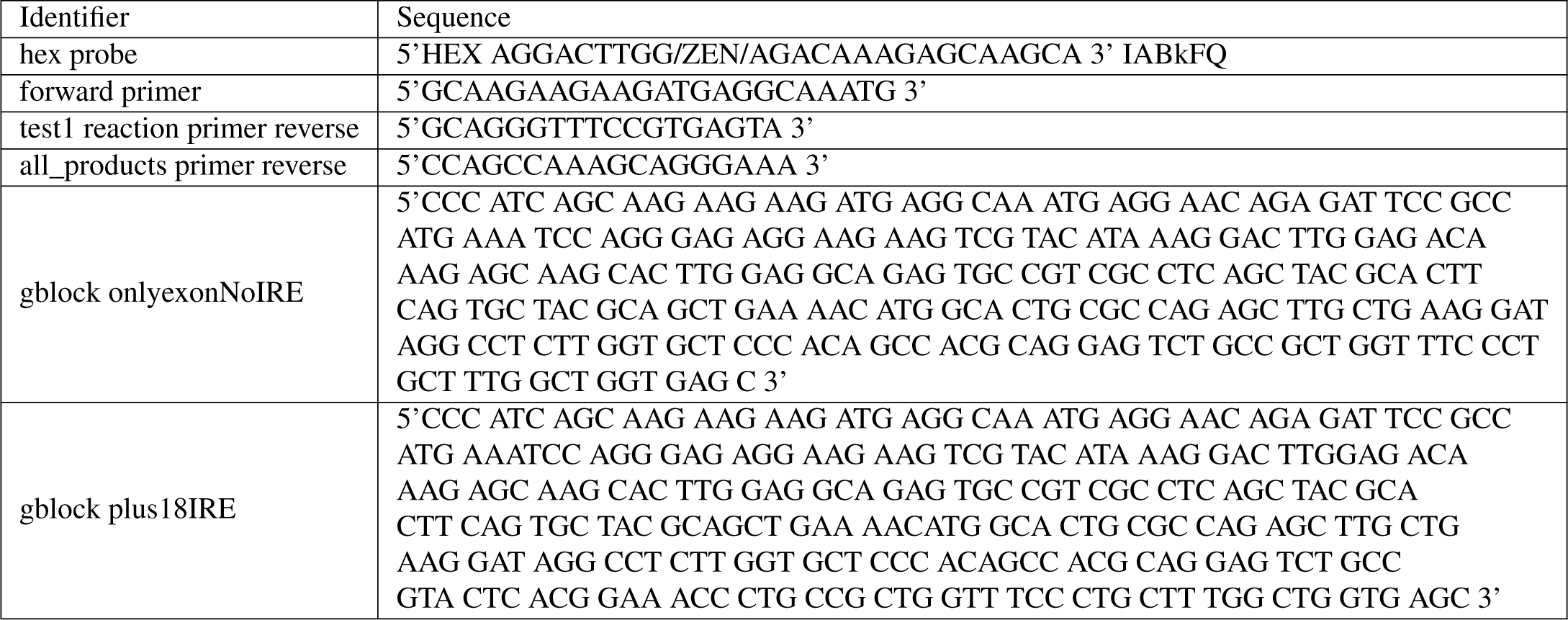
Primers and Probes

### Gene Expression Data Analysis

The amount of HEX fluorescence in each cycle (Supplemental Results File 1) was fit to a five-parameter curve using the qPCR R package, https://CRAN.R-project.org/package=qpcR^24^. Code for the fit is available at https://github.com/tbillman/Stapleton-Lab/blob/master/qpcR/3-7-18/3-7-18qPCRAnalysis.rmd. To calibrate the realtime assays we used synthesized DNA sequences (gBlocks), one with the small intron sequence included and the other with the IRE1-spliced-out sequence (Table 1). The two realtime reactions for each sample were adjusted to compensate for different reaction efficiencies using the positive control synthetic DNA reactions, as described at https://github.com/michael-byrd/Stapleton_Lab/blob/development/qPCR/adjustment/README.md. The Cp values calculated from the qPCR R package were changed to units of femtograms by using the Cp values from the femtograms of synthesized DNA in the calibration reactions (Supplemental Results Files 3 to 5). Sample replicates with no detectable realtime signal (i.e. with a Cp value of 40 for either of the two reactions) were replaced with the median-interpolated value once per replicate set.

### Decision Tree Recursive Partitioning Data Analysis

Decision tree partitioning is a nonparametric approach which uses the logworth P value calculation to find minimum sums of squares ‘splits’ for measurements using experimental factor levels (https://www.jmp.com/support/help/14/partition-models.shtml#). We analyzed the measured IRE-splicing difference values (after slope efficiency adjustment, interpolation and unit conversion to femtograms) using the partitioning tools in JMPv13 (SAS, Inc., Cary, NC). Splits were made until the minimum RMSE and maximum RSquared were achieved. The splits were visualized using the graphical tree widget in Orange (Demšar et al., 2013).

### Generalized Regression Data Analysis

The overall distribution for each response trait was evaluated in JMP statistical software to determine which distribution best fit the data. From a list of all tested distributions, we selected the distribution with the lowest AICc which was enabled within the generalized regression module in JMP. Once an appropriate distribution was selected a full factorial analysis using temperature and hormone treatment was fit to each phenotype with LASSO shrinkage based on the minimum AICc. Model fit details with full P-value listing and effect estimates are available in Supplemental Results File 6.

## Data Availability

All raw and processed data files and metadata are available in Figshare at DOI 10.6084/m9.figshare.7609931, https://figshare.com/s/175af4c449c8ccfe9a64.

## Results

A block design with plant growth regulator treatments applied within levels of growth temperature was used to determine the effect of treatment, temperature, and the interaction between treatment and temperature (Figure 1).

**Figure 1.**
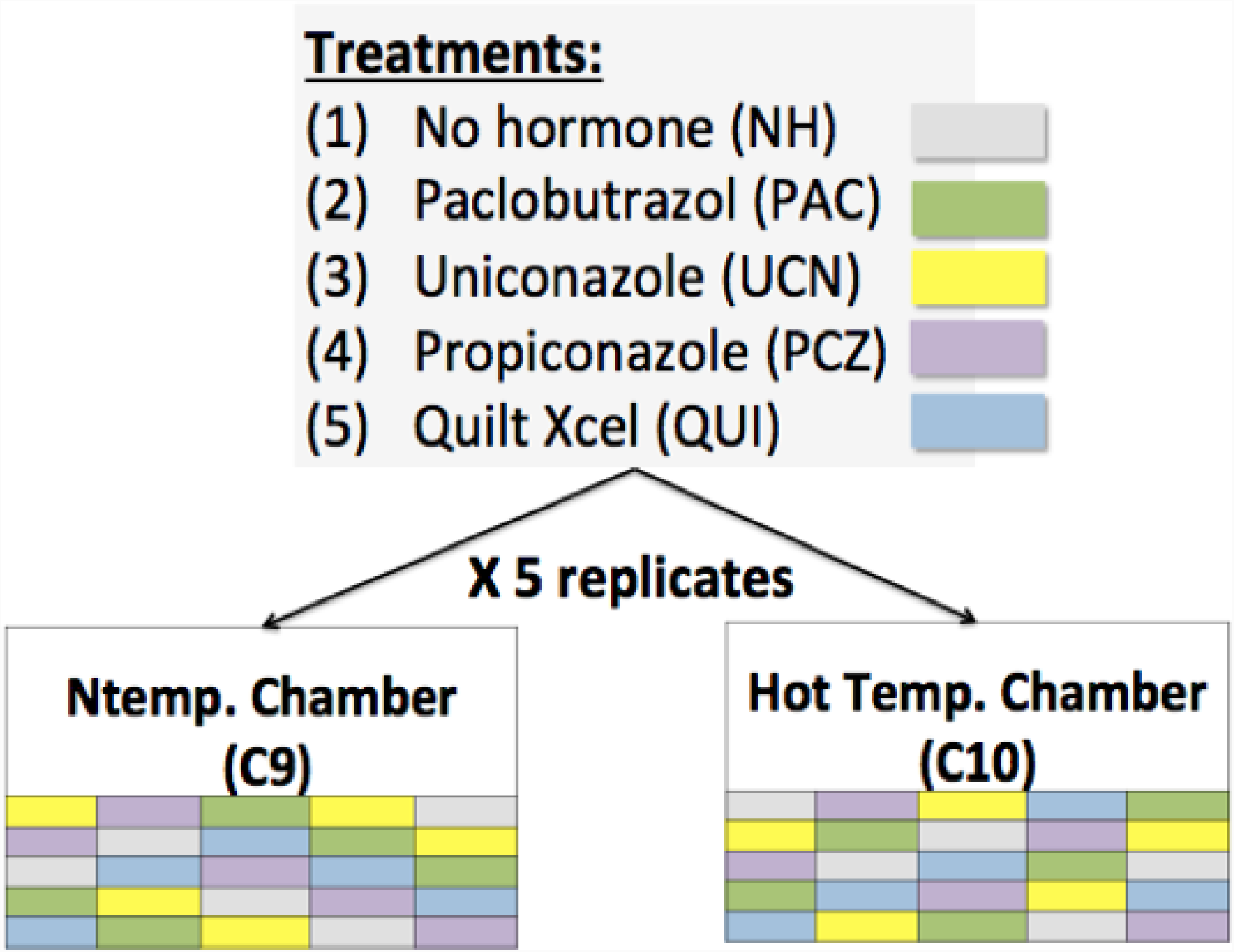
Experimental design layout, with designated chambers for heat stressed conditions and normal temperature conditions. Each treatment had five replicates in each chamber.

### IRE-UPR bZIP60 splicing effects

To measure the level of splicing of bZIP60 mRNA splicing we amplified the spliced and unspliced bZIP60 products, calculated the curve fit and inflection point for the amplification reaction over cycles, then took the difference between the calculated values to derive the amount of spliced product. The realtime qPCR assay was calibrated using serial dilutions of the synthesized DNA segments and the analysis included an adjustment for the difference in efficiency of amplification of the two reactions, to ensure that the product levels could be compared fairly across the full range of input DNA amounts (Supplemental Results Files 3,4,5).

Recursive partitioning was used to non-parametrically examine the contributions of temperature and treatment to the amount of IRE-induced splicing in bZIP60 messenger RNA from leaf tissue. The extent of the IRE1 UPR was partitioned by temperature and treatment factors that best split the dataset (Figure2). The first level split was between low no-hormone IRE UPR levels and higher plant growth regulator treatment levels (first level of the binary tree in Figure 2), with the non-hormone treatment having the lowest level of bZIP60 mRNA splicing (mean of 0.5 fg as compare to plant growth regulator mean of 1.4 fg). The temperature splits within plant growth regulator treatment levels (at level three of the three) show that high-temperature heat stress induced RNA splicing levels were always higher than the normal-temperature splicing response levels. Additional details on logworth values and model fit are provided in Supplemental Results File6.

**Figure 2.**
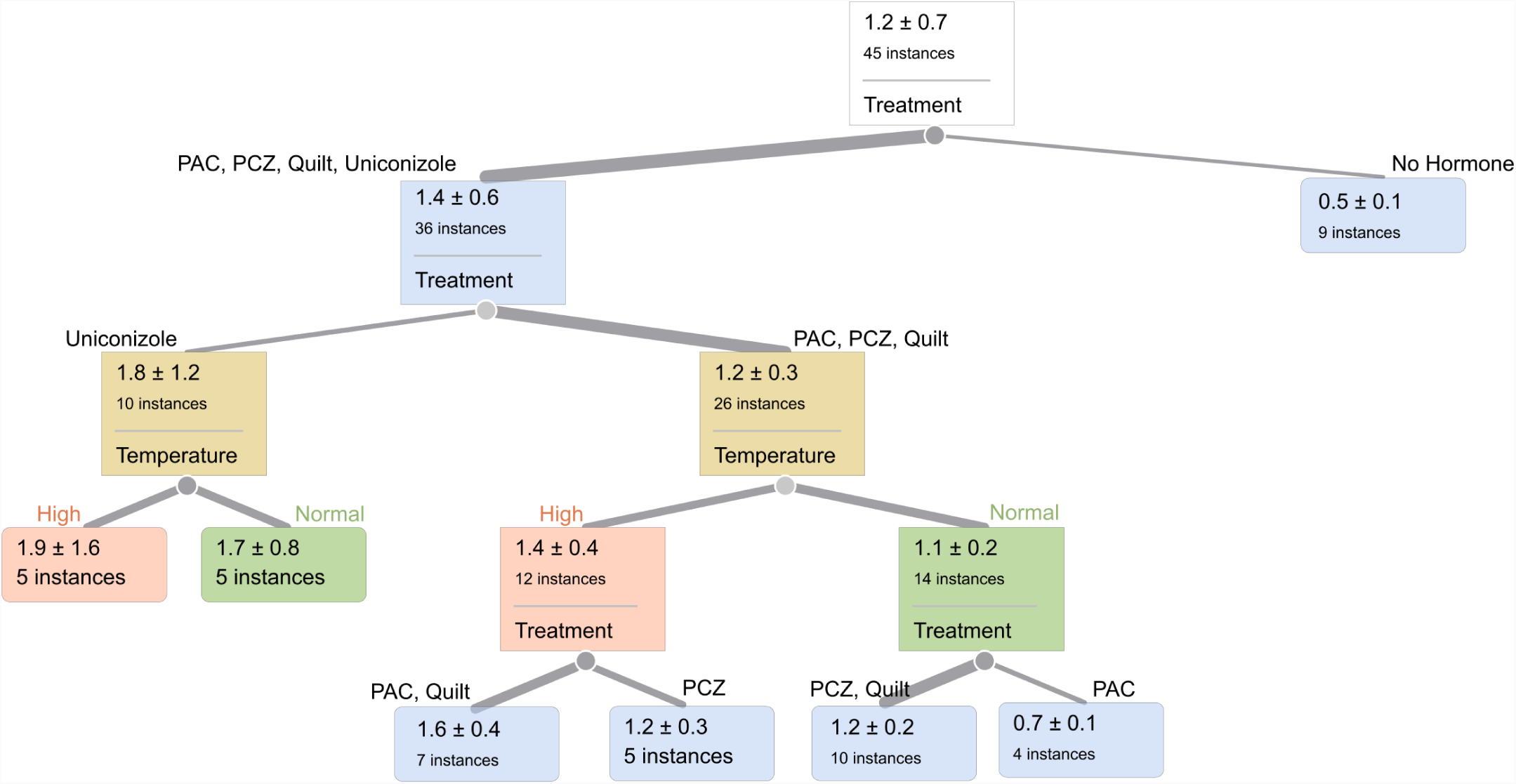
Visualization of IRE-specific bZIP60 splicing amounts (in femtogram, fg), with factor level partitions shown in order of importance from the entire data set (indicated in the top box). The RMSE for the tree was 0.675 and the RSquare was 0.309. Growth regulator treatments are non-hormone control, paclobutrazol (PAC), propiconazole (PCZ), Quilt, and uniconazole. The instance value is the n for that split, with mean (in femtograms, fg) and standard deviation (plus-minus) provided within the split details box at each level. The tree branch widths reflect the number of instances, with larger branch widths indicating larger n.

Generalized linear regression analysis with variable selection identified a significant effect of plant growth regulator treatment, with the overall P value for the treatment effect equal to 0.0002 (Figure3). When factors and factor interactions were analyzed using the Lasso model fit (Figure3), no-hormone splicing level was significantly lower than uniconazole (P<0.0001) and PAC significantly less than uniconazole (P=0.0404). Additional details on model fits are available in Supplemental Results File6.

The variance in treatments is visually different in the violin plots (Figure3), with uniconizale treatment showing a much larger range (as well as a larger mean and median than the no-hormone control). We tested for variance differences using Levene’s measure (Schultz, 1985) and found no significant differences in variance in pairwise comparisons (Supplemental Results File 1). However, a nonparametric analysis of variances indicated that the uniconazole treatment variance was significantly higher than the overall variance (Supplemental Results File6). The Levene’s test assumes that the groups are balanced and the noise is symmetric, while the non-parametric test does not have the symmetric noise assumption.

To test for differences in temperature within plant growth regulator treatments, as visualized in the decision tree (Figure2), we fit a generalized regression with the temperature factor considered within each plant growth regulator treatment. There was a significantly (P=0.0075) higher level of heat stress induced splicing in the PAC treatment high-temperature sample (Figure4). Plant growth regulator and growth temperature thus interact, with significant temperature effects seen in the paclobutrazol treatment (Figure4) and a trend toward higher splicing in elevated temperature in the no-hormone treatment (P=0.08, Supplemental Results File6).

### Plant growth effects

Plant heights were significantly reduced by plant growth regulator treatment, with P<0.0001 (Figure5). Factor comparisons significant in the model fit included the no-hormone treatment, which was significantly higher than uniconazole (P<0.0001) and the PAC treatment, also significantly higher than uniconazole (P=0.0002).

The interaction between temperature and treatment for plant height was significant with P=0.0425. Specific factor combinations that were significant included the comparison between elevated temperature and normal heights in the Quilt treatment and between high and normal temperatures in uniconazole (P=0.0025), with increased height in Quilt high temperature plants and shorter plants in uniconazole high compared to normal temperatures (Figure6).

Plant above-ground biomass and root biomass were also decreased in the plant growth regulator treatments compared to no-hormone (Supplemental Results File6 section B). The PAC treatment plants exhibited significantly lower root mass in elevated temperatures (Supplemental Results File6).

## Discussion

In this work we addressed the urgent outstanding question of whether plant growth regulators can influence the plant UPR. We found that under elevated temperature PAC-treated plants exhibited an increased bZIP60 mRNA splicing. Thus, we propose that PAC application ‘primed’ the plants, generating a stronger UPR response to increased heat. These results may have practical implications in understanding how plant growth regulators may facilitate growth through the modulation of specific stress responses.

In the partition analysis of bZIP60 mRNA splicing levels, the PAC and Quilt group exhibit a larger response in high temperature (Figure2) than PCZ. Previous studies of paclobutrazol in combination with another triazole in maize seedlings indicated that the combination of two triazoles increased tolerance to a high-temperature stress^25^, suggesting that the exact dose of plant growth regulator may be important. Quilt contains PCZ, which is known to affect brassinolide pathways^26^; these two growth regulators have similar response patterns in our study, with a slightly higher RNA splicing levels than the control and very little difference in splicing ratio at high temperature compared to normal temperature (Figure2). This suggests that the PAC effects could be mediated through its effects on the gibberellic acid pathway instead of effects on the brassinosteroid pathway. This hypothesis could be tested in genetically tractable models using mutants, or by time series and chemical interventions to capture the order and type of activation of particular pathway components.

Uniconizole and PAC have similar modes of action in the gibberellic acid pathway, though uniconazole also affects zeatin pathways (Best et al., 2017), suggesting that zeatin might suppress the UPR effect of PAC or that the uniconazole effect on gibberellic acid pathways is qualitatively different than the PAC effect. Uniconizole effects in maize do depend on the applied dose^21^, and our concentration was higher than the optimal amount determined by Ahmad et al. (2018), so we might expect to see a negative effect of uniconizole treatment in our experiment. The uniconazole treatment exhibited high variance (Figure 3, Supplemental Results File6), which suggests that the regulatory circuitry for the UPR has multiple, possibly competing, inputs from uniconizole^27^. Typically high variance is seen when there are two or more distinct activities from an input; variance mapping of the uniconizole response could thus be expected isolate pathway components^28^.

**Figure 3.**
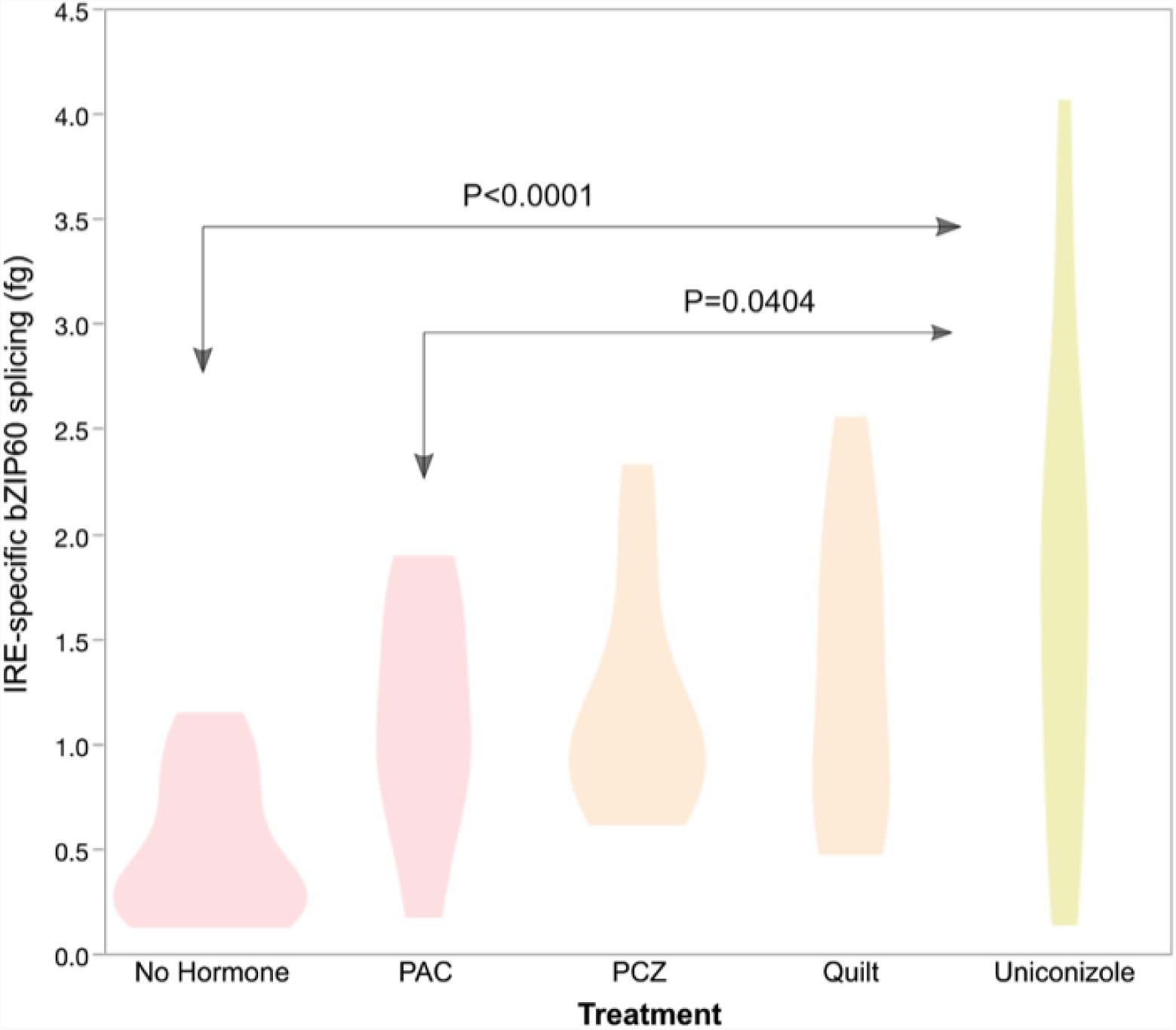
Probability density plot of IRE-specific bZIP60 splicing (in femtogram, fg) with significant comparisons indicated by pale red and yellow colors and arrows; beige treatment violins were not significantly different from the pale red or yellow treatments. The probability density indicates the data point range, with wider areas indicating values more likely to be found in the data. Growth regulator treatments are non-hormone control, paclobutrazol (PAC), propiconazole (PCZ), Quilt, and uniconazole.

**Figure 4.**
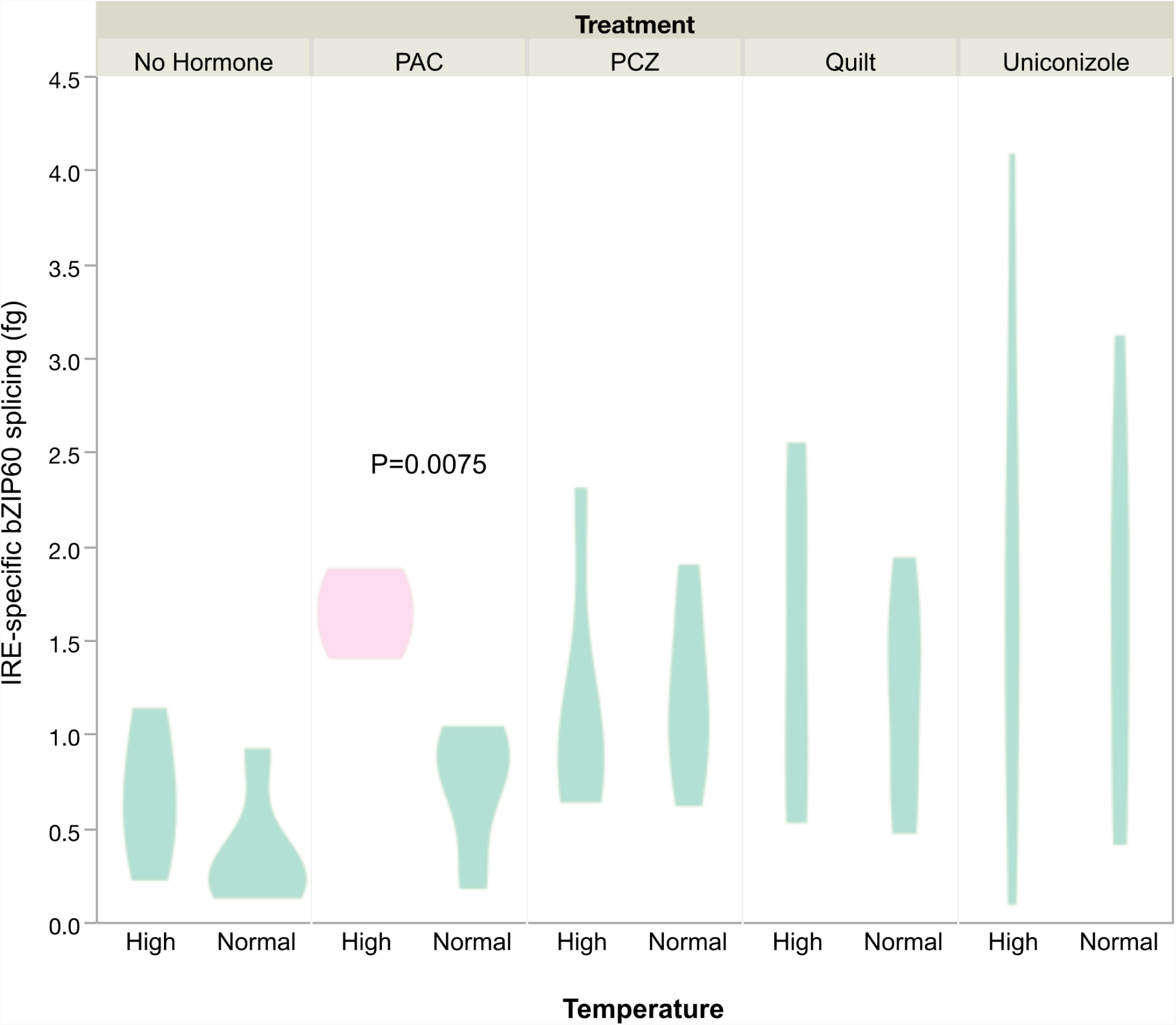
Probability density plot of IRE-specific bzip60 splicing (in fg) with significant comparisons indicated by different colors. Higher values on the y axis indicate higher levels of IRE1-specific splicing. Growth regulator treatments are non-hormone control, paclobutrazol (PAC), propiconazole (PCZ), Quilt, and uniconazole.

**Figure 5.**
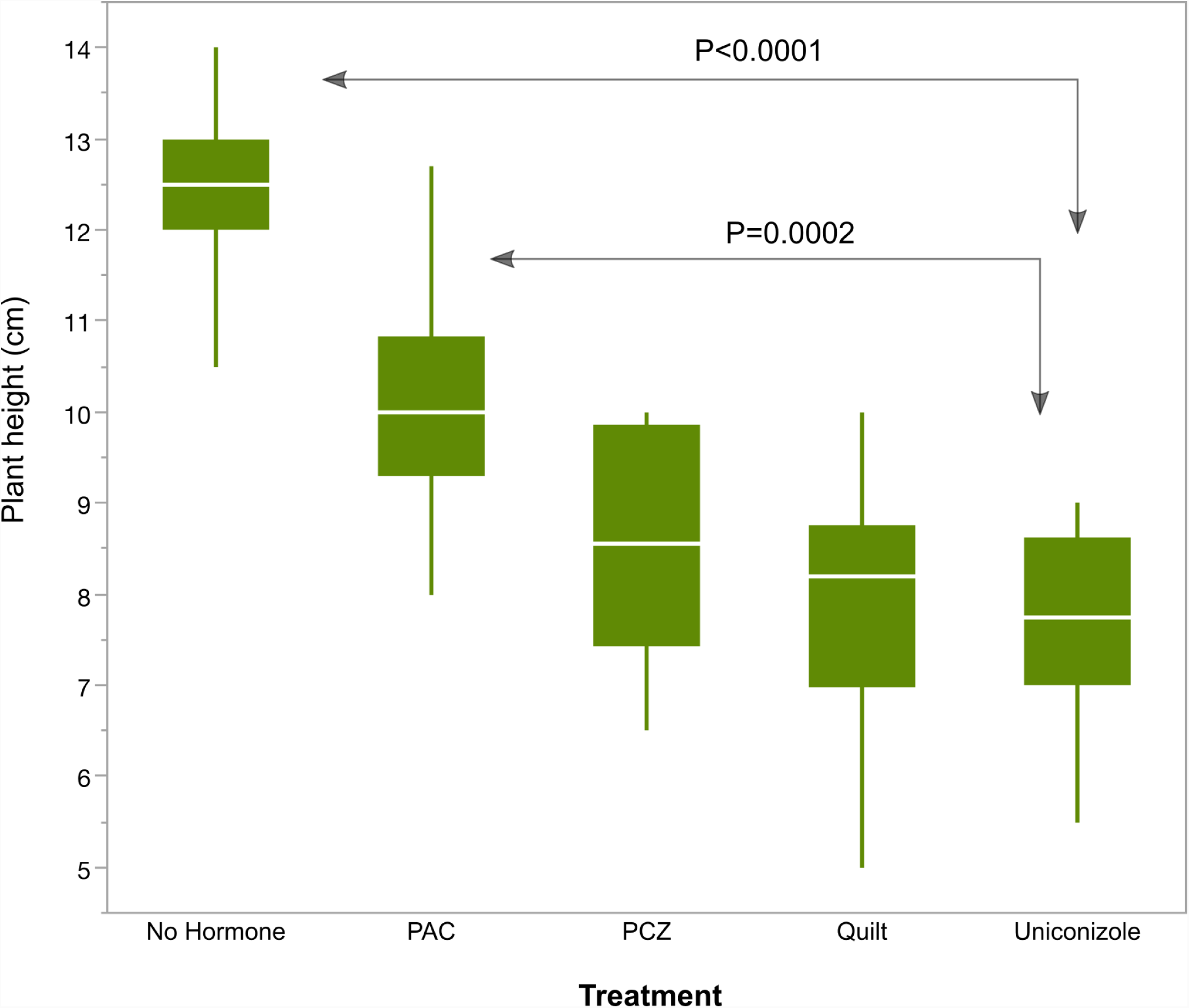
Box plot of plant heights (in cm). Means are indicated with white horizontal bars. Growth regulator treatments are non-hormone control, paclobutrazol (PAC), propiconazole (PCZ), Quilt, and uniconazole.

**Figure 6.**
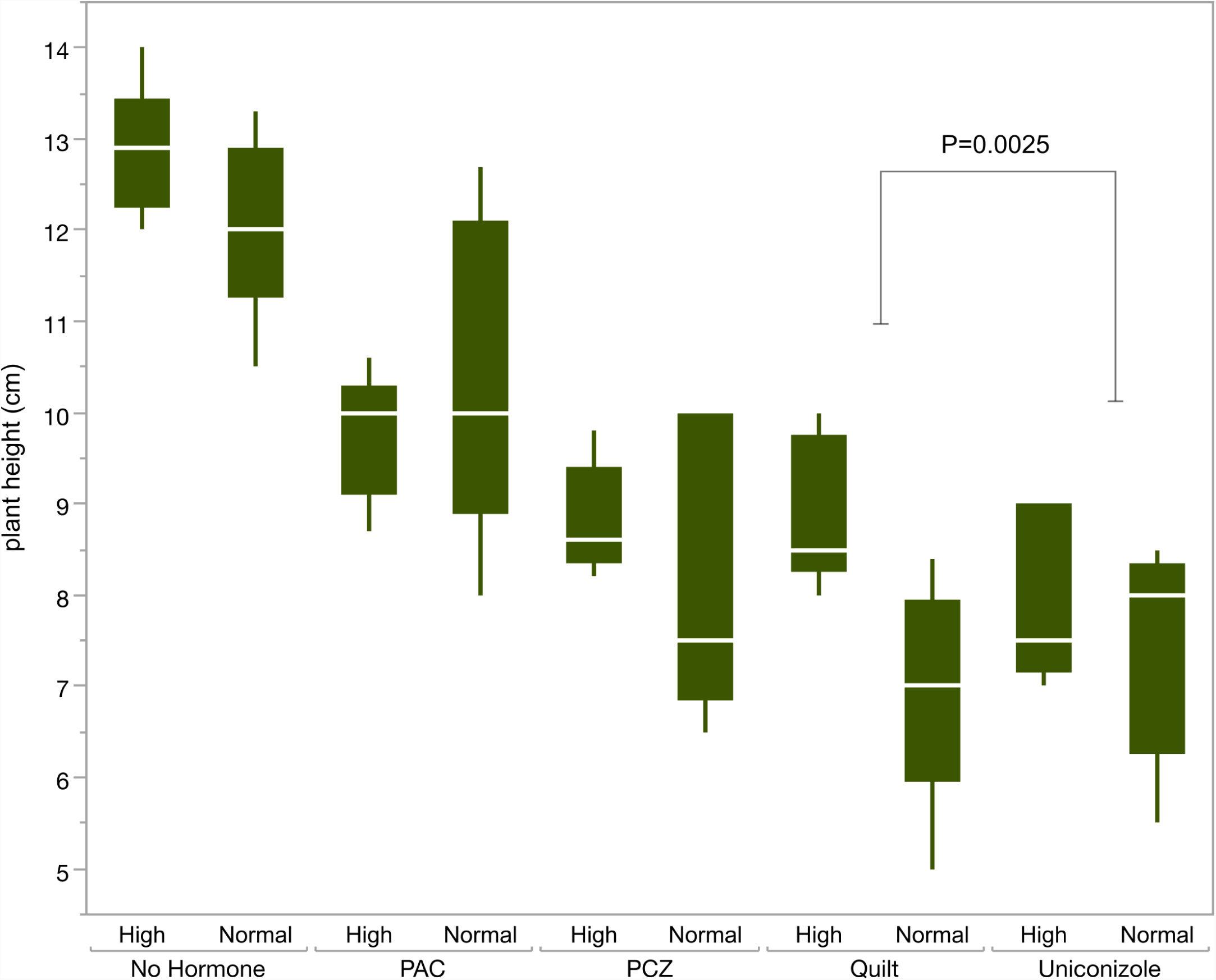
Box plot of plant heights (in cm) for each level of treatment and temperature. Means are indicated with white horizontal bars. Growth regulator treatments are non-hormone control, paclobutrazol (PAC), propiconazole (PCZ), Quilt, and uniconazole.

As expected, plant height was always lower in plant growth regulator treatment conditions (Figure5). This may be at least partially linked to activation of the UPR, as the UPR is known to participate in plant growth control^29^. However, we would need interventional experiments such as over-expression and under-expression of bZIP60 to determine the order of events between growth-regulator-induced plant height reduction and UPR signaling steps.

In conclusion, we demonstrate that paclobutrazol affects heat induced UPR signaling in maize, which is the first link between non-PIF4 and PIF4-related thermosensory signaling in plants and addresses a key question identified by Li (2018). If the PAC effect is through its inhibition of gibberellic acid biosynthesis, we would hypothesize that DELLA mutants would be expected to reduce bZIP60 mRNA splicing in response to heat stress. Similar hypotheses can be constructed for the other pathways affected by PAC. Combinations of mutants and chemical treatments are recommended for future analysis of these pathways^30^. Our study has provided new evidence that plant growth regulators may act on heat stress signaling via the UPR.

## Acknowledgements

We appreciate assistance with maize GDb bZIP60 sequence annotation from Iowa State University postdoctoral associate Zhaoxia Li and contributions to the manuscript text and organization from Dr. Stephen Howell, ISU. We thank the NCSU Phytotron staff for their technical assistance and expert care of the plants and growth chambers. This material is based upon work supported by the National Science Foundation under Grant No. 1444339. Any opinions, findings, and conclusions or recommendations expressed in this material are those of the author(s) and do not necessarily reflect the views of the National Science Foundation.

## Author contributions statement

E.N. and A.E.S. conceived the experiment, E. N. conducted the experiment, M.S.B., T. B. and A.E. S. analyzed the data, F.B and A.E.S. wrote the manuscript. All authors reviewed the manuscript.

## Additional information

The authors declare no competing interests.

